# A new chemical carcinogen and Western diet protocol to reliably induce advanced hepatocellular carcinoma in male and female mice

**DOI:** 10.64898/2026.03.06.710198

**Authors:** Melina C. Mancini, Elise R. Adams, David H. Burk, Richard Carmouche, Sara Webb, Jaroslaw Staszkiewicz, J. Michael Salbaum, Samuel G. Mackintosh, Timothy D. Heden

**Affiliations:** Molecular Metabolism Lab, Pennington Biomedical Research Center, Baton Rouge, LA, U.S.A.; Cell Biology and Bioimaging Core, Pennington Biomedical Research Center, Baton Rouge, LA, U.S.A.; Genomics Core, Pennington Biomedical Research Center, Baton Rouge, LA, U.S.A.; Molecular Mechanisms Core, Pennington Biomedical Research Center, Baton Rouge, LA, U.S.A.; Department of Biochemistry and Molecular Biology, University of Arkansas for Medical Sciences, Little Rock, AR, U.S.A.

**Keywords:** Hepatocellular carcinoma, chemical method to induce liver cancer, liver spatial transcriptomics, proteomics, diethylnitrosamine, thioacetamide, sucrose

## Abstract

**Background and Aims:** A combination of chemical carcinogens and Western diet have been used to induce hepatocellular carcinoma (HCC) in mice, but these models show low incidence of HCC in female mice, while mice that do develop HCC show a wide range of HCC stage development. Therefore, a more reliable mouse model that induces advanced stage HCC in both male and female mice is warranted, which will enable reliable preclinical testing of therapeutics for advanced HCC. The purpose of this study was to create a simple, yet tolerable and reliable chemical carcinogen and Western diet induced mouse model of advanced stage HCC for both male and female mice.

**Approach and Results:** We report that providing mice with a Western diet at weaning and for their lifetime combined with the sequential administration of low dose diethylnitrosamine (DEN), thioacetamide (TAA), and sucrose water promotes stage 2-3 HCC by 30 weeks of age in 100% of male mice and 96% of female mice (4% had stage 1 HCC), with a low fatality rate. Spatial transcriptomics, proteomics, and western blotting revealed this model alters genes and proteins similar to human HCC.

**Conclusions:** This study introduces, for the first time, a reliable chemical carcinogen model that induces advanced HCC in both male and female mice. Importantly, the model was tolerable for mice and induced protein and gene signatures comparable to human HCC. This new protocol will be a valuable model for preclinical testing of new therapeutic approaches for advanced HCC.

## Introduction

Hepatocellular carcinoma (HCC) is the most common liver malignancy (> 90% of cases) (1) and the fifth most common cancer worldwide (2). More than 800,000 people are diagnosed with HCC worldwide each year (1). The five-year survival of untreated HCC is 9.6% (3). While treatments such as liver transplantation, tumor ablation, open resection, minimally-invasive resection, transarterial (chemo)embolization, or selective internal radiation therapy can improve five-year survival rates, the majority of HCC patients do not have access to these therapies (3). Thus, more research is needed to help develop more practical and far-reaching treatment strategies for HCC.

The development of treatments begins with efficacy studies in preclinical models. In mice, diethylnitrosamine (DEN) is a common genotoxic carcinogen given to promote HCC initiation. DEN reacts with cytochrome P450 enzymes in the liver to form alkylating metabolites that alkylate deoxyribonucleic acid (DNA), forming mutagenic DNA adducts that promote HCC. The traditional approach includes administering a single dose of DEN via an intraperitoneal injection to young mice at two weeks of age. When mice are fed standard chow diets, this approach is well tolerated, but it can take 10 months (∼43 weeks) or longer to promote HCC (4-9). Moreover, there is an uneven occurrence time with some reports indicating that not all mice develop HCC (6, 7, 9, 10), with female mice having a much lower incidence compared to male mice (11-13). Combining DEN with other chemotoxic agents such as thioacetamide (TAA), carbon tetrachloride, and/or a Western diet (9, 14, 15) has been shown to increase the occurrence and speed at which HCC develops, but female mice still have less HCC occurrence, with some mice not developing HCC even at 40 weeks of age (15). Moreover, given the fact that HCC is most often diagnosed in humans at an advanced stage (3), a protocol is warranted that can consistently induce advanced stage HCC in both male and female mice, which would be a useful tool to not only study the pathogenesis of HCC but allow the reliable testing of new therapeutics.

Here we report a tolerable and reliable method to induce advanced stage 2-3 HCC in both male and female mice. The protocol involves providing mice with a Western diet at weaning and for their lifetime combined with multiple low dose DEN injections starting at 2 weeks of age for eight weeks, followed by TAA drinking water for four weeks, and 10% sucrose water for the remainder of their life. At 30 weeks of age 100% of male mice and 96% of female mice develop advanced stage 2-3 HCC with large visible tumors that have a gene and protein expression pattern comparable to human HCC.

## Methods

### Mouse Model

This study was approved by the Pennington Biomedical Research Center’s Institutional Animal Care and Use Committee. Mice of the strain C57BL6 were used for this study. Two groups of mice were used: 1) a low-fat diet (LFD) control group (Inotiv TD.05230, 18.7% protein, 68.7% carbohydrate, 12.6% fat), and 2) a Western diet (WD) fed group (Inotiv, TD.88137, 15.2% protein, 42.7% carbohydrate (34% sucrose), 42% fat) that received the chemical carcinogens diethylnitrosamine (DEN) and thioacetamide (TAA) (WD+DEN/TAA). Mice were placed on their respective diet after weaning. Mice in the WD+DEN/TAA group began receiving intraperitoneal injections of DEN once a week for eight weeks starting at two weeks of age (**Figure 1A**). This included 20 mg of DEN per kg of body weight for the first week, 30 mg of DEN per kg of body weight the second week, and 50 mg of DEN per kg of body weight for the next six weeks. This protocol was modified from previous reports (16, 17). One week after completion of DEN injections, mice were provided with TAA drinking water (300 mg/L) for a period of four weeks straight to promote liver damage, liver fibrosis, and accelerate HCC development. After two weeks of regular water, mice received 10% sucrose water to further promote HCC (9, 15), and the sucrose water was provided until euthanasia at 30 weeks of age. All animals were group housed with a 12-hour light and 12-hour dark cycle at room temperatures ranging between ∼22-23°C. All mice were euthanized at 30 weeks of age. Two mice (out of 56 mice, 3.5%) in the cancer group were euthanized prior to 30 weeks of age due to poor physical condition, indicating that the protocol is highly tolerable.

**Figure 1.**
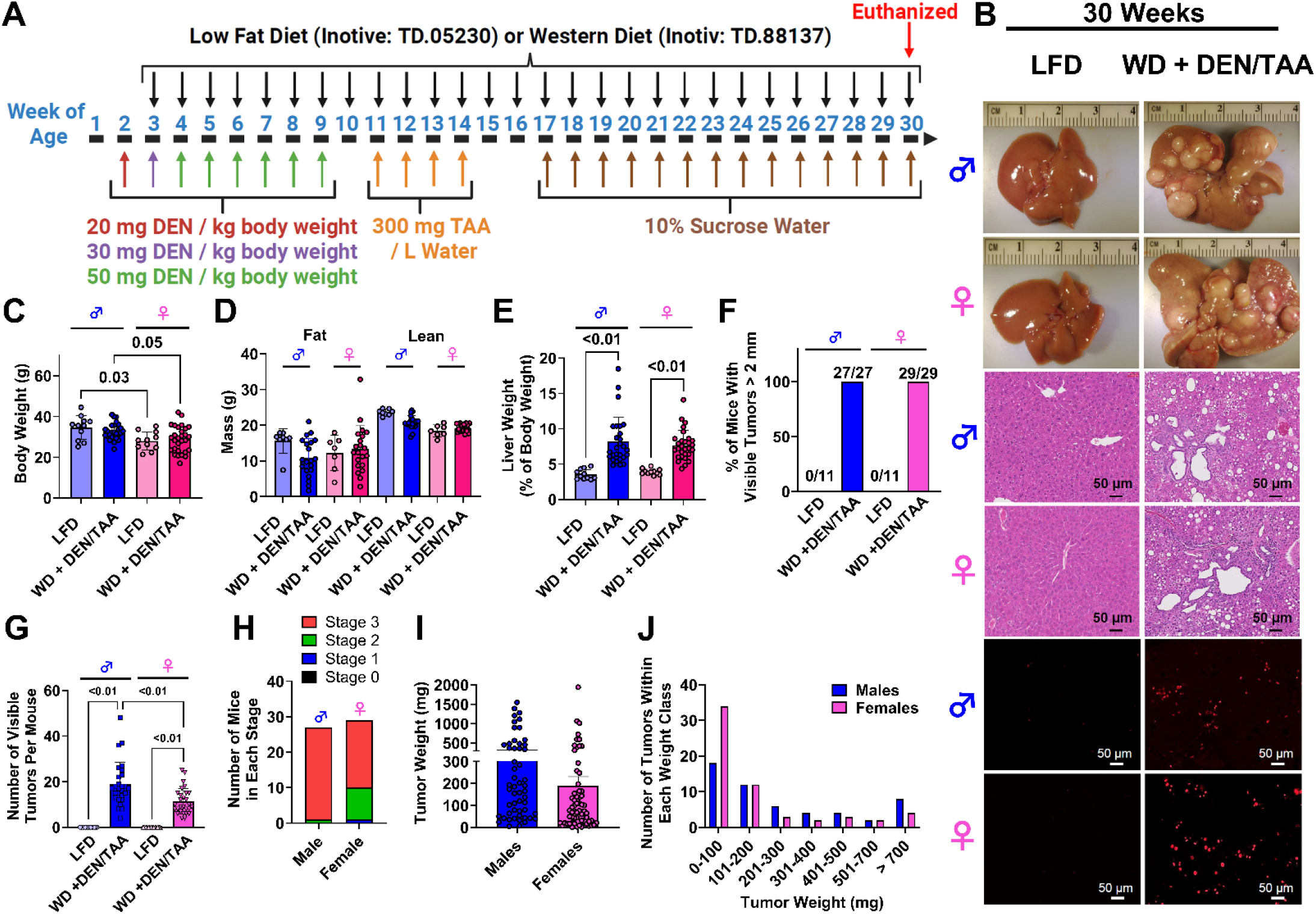
A protocol to reliably produce advanced stage 2-3 HCC in male and female mice. **A)** Illustration showing protocol details to induce advanced HCC in male and female mice. The protocol involved the sequential administration of DEN, TAA, and 10% sucrose water, all on top of a Western diet. **B)** Representative image of whole liver (top), hematoxylin and eosin staining (H&E) (middle), and Ki-67 staining (bottom) of liver sections from male and female mice. **C)** Final body weight of the mice. **D)** NMR measured body composition including fat mass and lean mass. **E)** Liver weight, normalized to body weight. **F)** Percentage of mice with visible tumors > 2 mm in diameter. **G)** The total number of visible tumors per mouse. **H)** Number of mice with stage 0, stage 1, stage 2, or stage 3 HCC. **I)** Tumor weights. A total of 3 tumors per mouse were weighed and included in the data. **J)** Frequency distribution of a range of tumor weights. Where applicable, data are presented as means ± S.E.M.

Prior to euthanasia, mice were anesthetized with a cocktail of ketamine (120 mg per kg of body weight), xylazine (9 mg per kg of body weight), and acepromazine (2 mg per kg of body weight) in saline via intraperitoneal injection. Once deep anesthesia was reached, blood was collected through a cardiac blood draw. Tissues were harvested after cervical dislocation. Tissues and serum were stored at −80°C.

### Liver Imaging

Once the liver was removed, a picture of the liver was taken using a digital microscope camera (Opqpq), the liver was weighed, the total number of visible tumors were counted, and some tumor and adjacent, non-tumor tissues were dissected, weighed, and snap frozen in liquid nitrogen. Sections of liver tissue were preserved in 10% formalin overnight and then transferred to 70% ethanol prior to embedding and sectioning. Hematoxylin and Eosin (H & E) staining was performed on liver sections within the Cell Biology and Bioimaging Subcore of the Molecular Mechanisms Core at Pennington Biomedical Research Center. Standard approaches were used for staining, as previously described (17-20). Ki-67-staining was performed using the BOND RX Fully automated research stainer (Deer Park, IL, USA) with casein blocking solution (37583, ThermoFisher Scientific), the primary Ki-67 antibody at a concentration of 1:300 (ab15580, abcam), and a goat anti-rabbit Alexa Fluor 647 secondary antibody (A32733, Invitrogen, Waltham, MA, USA) at a 1:300 dilution as previously described by our lab (17). Slides were scanned using a Zeiss Axioscan 7 slide scanner.

### Body Weight and Composition

The body weight of the mice was measured using a digital pet scale for small animals (Mindpet-med). The body composition of the mice was measured using a Minispec LF90 Series TD-NMR System.

### 10X Visium Spatial Transcriptomics

Spatial transcriptomics was performed within both the Cell Biology and Bioimaging Core and the Genomics Core at Pennington Biomedical Research Center. Formalin fixed paraffin embedded (FFPE) tissue sections were used for spatial transcriptomics, which were performed according to the 10X Genomics Visium Spatial Gene Expression Reagent Kit. Tissues were sectioned to 5 µm thickness and the slice was placed on the Visium Spatial Gene Expression Slide with barcoded spots. Next, paraffin was removed and the tissue was permeabilized to release mRNA. Reverse transcription was performed using the 10X Genomics Visium Spatial Tissue Optimization Kit, relying on oligo-dt primers seeded in 55 µm diameter capture spots on the Visium slide. Primers contain a spatial barcode to permit localization of sequence reads and unique molecular identifiers to exclude PCR-based duplicate reads. Lastly, the library was prepared using the 10X Genomics Spatial Gene Expression V2 Kit. Sequencing was performed using an Illumina NextSeq.

### Serum Aspartate Aminotransferase (AST)

Serum aspartate aminotransferase (AST) activity was measured using a commercially available assay kit (NBP3-24470, Novus Biologicals, LLC, Centennial, CO, USA), according to manufacturer specifications.

### Serum Alanine Aminotransferase (ALT)

A commercially available assay kit (E-BC-K235-M, MSE Supplies LLC, Tucson, AZ, USA) was used to measure serum ALT activity, as we previously described (17).

### Serum Major Urinary Protein 20 (MUP20)

A commercially available MUP20 ELISA kit (EKF58167, Biomatik, Ontario, Canada) was used to measure serum MUP20. Briefly, the ELISA plate was washed twice followed by being loaded with standards or samples. A biotin labeled antibody was added to each well, the plate was covered, and incubated at 37°C for 45 min. Next, the plate was washed three times, an HRP-Streptavidin Conjugate was added to each well, and the plate was incubated at 37°C for 30 min. Next, the plate was washed five times, a TMB substrate was added, and the plate was incubated at 37°C for 20 min. Immediately following the incubation, stop solution was added and the absorbance of the plate was measured at 450 nm.

### Western Blotting

Western blotting was performed as previously described (17-21). Liver samples were put into RIPA buffer (89900, Thermo Fisher Scientific, Plaquemine, LA, USA) with added phosphatase (A32957, Thermo Fisher Scientific) and protease (A32963, ThermoFisher Scientific) inhibitors and homogenized using a Teflon pestle. Homogenized samples were spun at 12,000 g for 10 min, the remaining supernatant was transferred to a new tube, and the total protein of the lysate was measured using a Pierce^™^ BCA Protein Assay Kit (23225, Thermo Fisher Scientific). Between 20-30 μg of protein was separated by SDS-PAGE (Bio-Rad Laboratories, Hercules, CA, USA), transferred onto PVDF membranes, and the protein content in the membrane was determined using Ponceau S staining solution (40000279, Thermo Fisher Scientific). The membranes were blocked in 5% TBST for one hour prior to being exposed to the primary antibody in 5% TBST overnight at 4°C. The primary antibodies included fatty acid binding protein 5 (FABP5, 39926T, Cell Signaling Technologies, Danvers, MA, USA), glycogen phosphorylase brain isoform (PYGB, 12075-1-AP, Proteintech, Rosemont, IL, USA), and mechanistic target of rapamycin (mTOR, total mTOR antibody: 2983S; Phosphorylated at site serine 2448 mTOR antibody: 2971S, both antibodies from Cell Signaling Technologies). After incubating with the primary antibody, the membranes were washed 5 times with TBST and then incubated with the anti-rabbit IgG, HRP-linked secondary antibody (7074, Cell Signaling Technologies) in 5% non-fat dry milk for 1 hour at room temperature. Next, the membranes were washed 5 times with TBST, chemiluminescent substrate (34578, SuperSignal EstPico Plus, Thermo Scientific) was added, and the membranes were imaged using an Odyssey Fc Imager (LI-COR Biosciences, Lincoln, NE, USA).

### Human Liver Biopsy Samples

De-identified human liver matched tumor and normal tissue lysates were purchased from Novus Biologicals (Centennial, CO, USA). These samples were supplied as a 1 mg/ml lysate solution that was directly used in Western blotting. Since the human samples were collected elsewhere and de-identified, the Institutional Review Board at Pennington Biomedical Research Center determined that this is considered “Not human subjects research” as defined by 45 CFR Part 46.

### Proteomics

The proteomics experiment and follow-up bioinformatics analysis were performed at the IDEA National Resource for Quantitative Proteomics within the Winthrop P. Rockefeller Cancer Institute at the University of Arkansas. Samples were prepped and proteomics were performed as has been described elsewhere(22, 23).

### Promethion Indirect Calorimetry

The Promethion Core (Sable Systems International) was used to measure oxygen consumption, respiratory exchange ratio, physical activity, and food intake. Approximately five days prior to metabolic testing, mice were initially placed in Promethion cages to acclimate mice to the new cage and living in isolation. Mice were tested for four consecutive days. On day three, mice were fasted for four hours prior to being injected intraperitoneally with 2 mg of total glucose (1.6 mg of ^12^C glucose and 0.4 mg of U^13^C glucose) per gram of body weight so that whole body glucose oxidation (^13^C carbon dioxide production) could be measured. Two hours after administering the glucose, mice were refed with their assigned diet.

### Statistics

Graphpad Prism 10.6.1 was used to perform some statistical analyses. A one-way ANOVA with follow-up Tukey’s comparisons were used to identify statistically significant effects (P < 0.05) between groups. Spatial transcriptomics were analyzed using Space Ranger to map reads to the spatial barcodes while Loupe Browser was used for visualization of the data. The spatial transcriptomics raw data was deposited into the Gene Expression Omnibus (GEO) data repository under series record GSE320362. ProteoDA was used to perform a statistical analysis on the proteomics data set. Proteins with an FDR adjusted P value < 0.05 and a fold change greater > 1.5 were considered significant. The Database for Annotation, Visualization, and Integrated Discovery (DAVID) was used to identify up-regulated or down-regulated cellular components and Kyoto Encyclopedia of Genes and Genomes (KEGG) pathways that were modified.

## Results

To establish a reliable protocol that induces advanced stage 2-3 HCC consistently in both male and female mice, we used the sequential administration of DEN, TAA, and 10% sucrose water while providing a solid Western diet from weaning age (**Figure 1A**). The protocol was highly effective and produced many large visible liver tumors in both male and female mice, which is consistent with advanced HCC development (**Figure 1B**, upper images). To further confirm the development of HCC, H&E was used and revealed clear cell carcinoma (large, irregularly shaped white (unstained) areas and a greater number of nuclei (blue stained areas) compared to LFD control mice (**Figure 1B**, middle images). Additionally, Ki-67 staining, a marker of cell proliferation that is upregulated in HCC, was highly prevalent in both male and female mice with HCC but was nearly undetectable in liver sections from LFD control mice (**Figure 1B**, lower images). Final body weight was lower in LFD or WD+DEN/TAA female mice compared to male mice (**Figure 1C**), with no impact of HCC on the final body weight within each sex. Consistent with this, there was no effect of HCC on fat or lean mass (**Figure 1D**). Liver weight was significantly elevated in both male and female WD+DEN/TAA groups compared to LFD control groups, indicating that large tumor mass increased the weight of the liver, an effect consistent with advanced HCC development (**Figure 1E**). 100% of male (27/27) and female (29/29) mice had visible tumors > 2 mm in diameter, while mice in the LFD groups had no visible tumors (**Figure 1F**). The total number of visible tumors was significantly greater in male mice compared to female mice (**Figure 1G**). We next classified the mice into HCC stages using a scoring approach previously described that uses the number of macroscopic liver tumors with a diameter > 2 mm as an indicator of HCC stage (24). Mice with no macroscopic liver tumors were considered stage 0, 1-3 tumors were considered stage 1 HCC, 4-6 tumors were considered stage 2 HCC, and 7 or more tumors were considered stage 3 HCC. Using this approach, 26 out of 27 male mice had stage 3 HCC, while 1 out of 27 had stage 2 HCC (**Figure 1H**). In female mice, 19 out of 29 had stage 3 HCC, 9 out of 29 had stage 2 HCC, and 1 out of 29 had stage 1 HCC. Tumor weight varied and had a large range in both male and female mice but was not significantly different between sexes (**Figure 1I-1J**). The majority of tumors that were weighed were between 1-100 mg for both sexes (**Figure 1J**). Collectively, these data indicate the protocol reliable produces advanced stage 2-3 HCC in both male and female mice.

To further confirm that HCC developed and determine if this protocol induces HCC that has a similar gene expression profile compared to human HCC, we employed spatial transcriptomics using the 10X Genomics Visium platform. There were a total of 2,282 spots under the tissue, with means of 39,147 reads and 1,389 genes per spot. Eosin staining revealed normal liver tissue areas as well as several areas of HCC (**Figure 2A**). The gene expression profile was classified into 4 clusters with unique gene signatures (**Figure 2A-2B, Supplemental Table 1**). Based on the morphological appearance of the tissue, cluster 1 and 2 are normal liver tissue areas. Cluster 1 (blue) had higher cytochrome P450 2e1 (Cyp2e1), ornithine aminotransferase (Oat), 6-phosphofructo-2-kinase/fructose-2,6-biphosphatase (Pfkfb1), and cytochrome P450 1a2 (Cyp1a2) cluster mean counts compared to cluster 2 (Supplemental Table 1), indicating these hepatocytes are likely perivenous hepatocytes as these genes have previously been validated to be more highly expressed within the perivenous zone (20, 25). Cluster 2 (orange) had higher Cdh1 cluster mean counts, which is a marker of periportal hepatocytes (20, 25), indicating this area is likely periportal hepatocytes. Cluster 3 (green) and cluster 4 (red) had gene signatures associated with HCC, albeit unique gene signatures highlighting that the tumor microenvironment can vary. Cluster 3 (green) and cluster 4 (red), compared to clusters 1 (blue) and 2 (orange), had higher abundance of markers known to be elevated in human HCC including fatty acid binding protein 5 (Fabp5) (26, 27) and lipocalin 2 (Lcn2) (28), indicating that the HCC produced from this protocol is similar to human HCC. Cluster 3 (green) had a higher cluster mean count of alpha fetoprotein (Afp), a marker of HCC (29), compared to cluster 4 (red), indicating that large tumors express Afp. Cluster 4 (red) had a high abundance of Hdlbp, a protein associated with HCC metastasis (30). A heatmap of the top 10 differentially expressed genes between each cluster illustrates the unique gene signatures of the different HCC clusters (**Figure 2C**). Overall, the WD+DEN/TAA protocol described here induced changes in gene expression that mirror some changes observed in human HCC.

**Figure 2.**
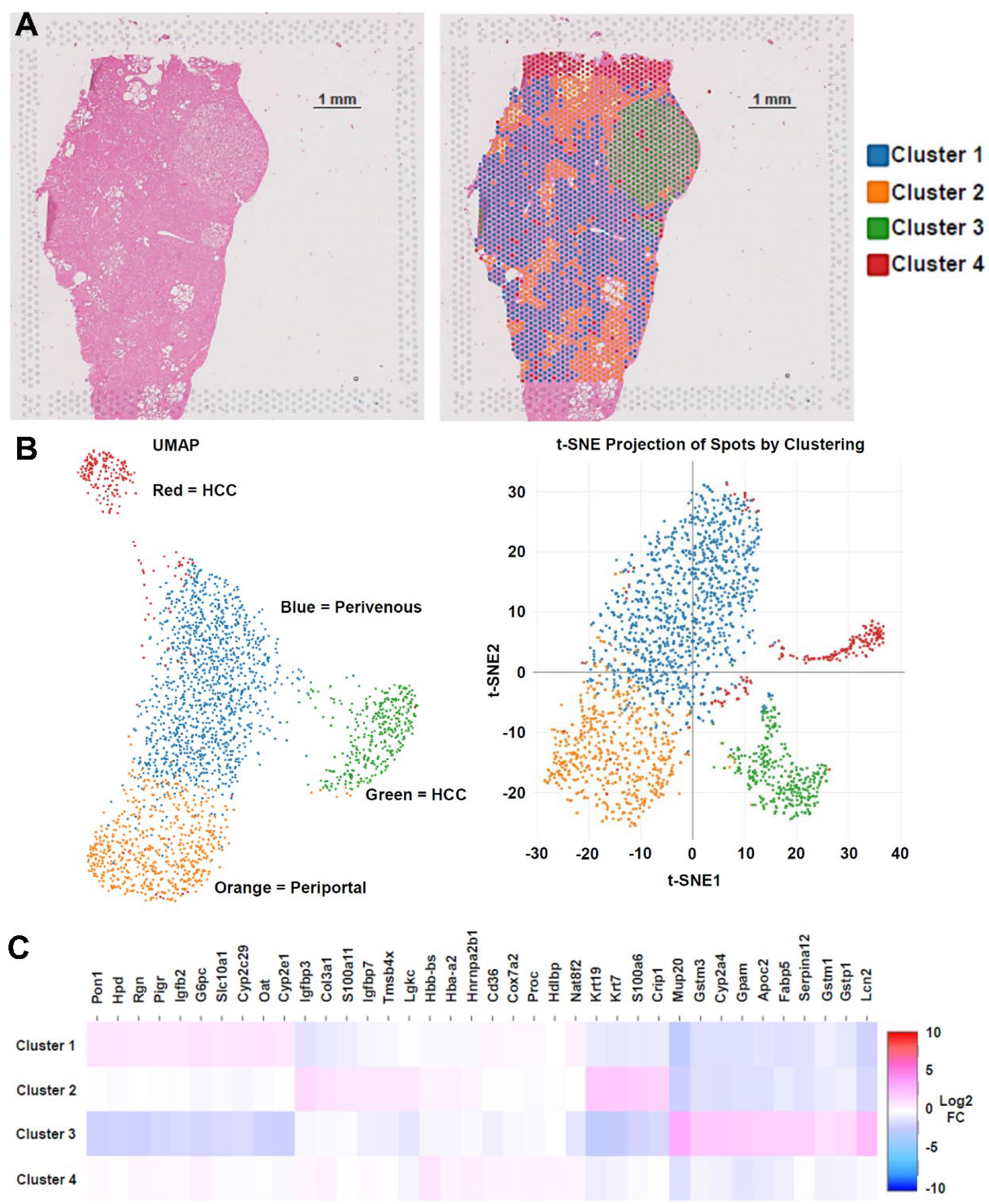
Spatial transcriptomics of mouse HCC. **A)** H&E image within the capture area on the Visium Spatial Gene Expression Slide slide (left) and H&E image covered with barcoded spots (right) that were clustered based on their gene signatures. **B)** A uniform manifold approximation and projection (UMAP) image visualizing the four gene clusters (left) and a t-distributed stochastic neighbor embedding (t-SNE) plot visualizing the four gene clusters identified in the liver tumor sample. **C)** Heatmap of select genes that were differential expressed within each cluster.

To further validate that this model induces advanced HCC, we also measured the protein abundance or activity of markers of HCC in the serum or liver tissue. Serum alanine aminotransferase (ALT) and aspartate aminotransferase (AST) are markers of liver damage and injury and are typically elevated in advanced HCC. Serum ALT and AST activity were both higher in the serum of WD+DEN/TAA treated male and female mice, compared to LFD male and female mice, providing evidence that advanced HCC has developed (**Figure 3A-3B**). The spatial transcriptomics data revealed an increase in major urinary protein 20 (MUP20) gene abundance in the liver tumor from male mice. Consistent with this, male mice had higher MUP20 serum protein levels compared to LFD fed mice (**Figure 3C**). However, LFD and WD+DEN/TAA female mice had similar serum MUP20 protein levels, indicating sex differences exist in this marker. Next, we performed western blotting on proteins known to be elevated in human HCC. The protein abundance of fatty acid binding protein 5 (FABP5) was significantly elevated in both mouse (**Figure 3D-3F**) and human (**Figure 3G**) tumor tissue compared to adjacent, non-tumor tissue (NT) where its abundance was low and barely detectable. Glycogen phosphorylase brain isoform (PYGB) was also elevated in mouse tissue compared to adjacent, non-tumor tissue. We also observed an increase in phosphorylation of the mechanistic target of rapamycin (mTOR) at serine 2448 (indicates higher mTOR activity) in mouse tumors compared to adjacent, non-tumor tissue and healthy tissue. Overall, these alterations in protein content are consistent with what has been observed in advanced human HCC.

**Figure 3.**
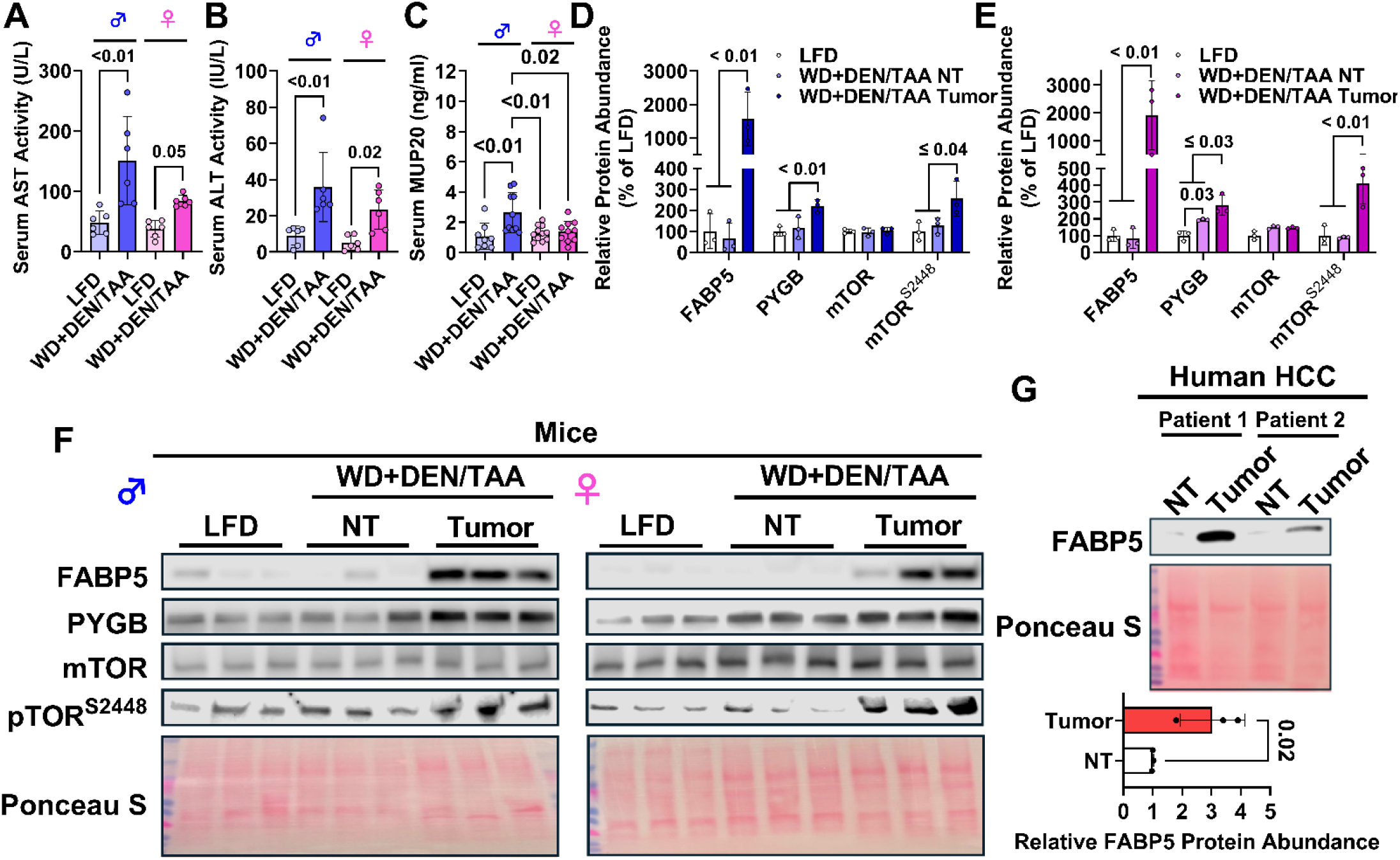
Markers of advanced HCC in serum and liver. **A)** Serum AST activity in LFD and WD+DEN/TAA treated male and female mice. **B)** Serum ALT activity in each group. **C)** Serum MUP20 levels in each group. **D)** Densitometry quantification of protein abundance in LFD, WD+DEN/TAA non-tumor tissue (NT), and WD+DEN/TAA tumor tissue treated male mice, from blot images shown in F. **E)** Densitometry quantification of protein abundance in LFD, WD+DEN/TAA NT, and WD+DEN/TAA tumor tissue treated female mice, from blot images shown in F. **F)** Western blot images of mouse liver proteins typically upregulated in HCC. **G)** Western blot images of human liver (NT and Tumor samples) proteins typically upregulated in HCC. Data are presented as means ± S.E.M.

Next, a proteomics analysis of tumor and adjacent, non-tumor tissue in WD+DEN/TAA treated mice was performed. A volcano plot revealed a total of 153 proteins were up-regulated and 161 proteins were down-regulated in tumor tissue compared to adjacent, non-tumor tissue (**Figure 4A, Supplemental Table 2**). FABP5 and PYGB were both up-regulated in the proteomics analysis, which is consistent with our western blotting data. Next, the DAVID Bioinformatics database was used to further characterize the proteome in this model. Functional annotation analysis revealed that the majority of up-regulated proteins were classified as endoplasmic reticulum or secreted cellular components (**Figure 4B, upper figure**), while the majority of down-regulated proteins were in the mitochondrion, endoplasmic reticulum, or microsome (**Figure 4C, upper figure**). A KEGG pathway analysis revealed that the majority of up-regulated proteins were within metabolic pathways or protein processing in the endoplasmic reticulum (**Figure 4B, lower figure**), while the majority of down regulated proteins were within metabolic pathways, steroid hormone biosynthesis, or arachidonic acid metabolism (**Figure 4C, lower figure**). A heatmap of the top 20 proteins (based on fold change) that were increased revealed FABP5 was elevated as well as proteins involved in terpenoid backbone biosynthesis including Hmgcr, Idi1, and Mvk (**Figure 4D**). A heatmap of the top 20 proteins (based on fold change) that were decreased revealed several proteins involved in steroid hormone biosynthesis and arachidonic acid metabolism including Cyp2c29, Cyp2c50, Cyp2c54, Cyp2e1, Cyp2c37, Hsd3b5, Srd5a1, Sult1w1, and Hpgd (**Figure 4E**). There were also 13 proteins that were only expressed in non-tumor tissue and not in tumor tissues, with three of these proteins being involved in glutathione metabolism including Nat8fs; Nat8f7, and Nat8 (**Figure 4F**). Conversely, 31 proteins were uniquely expressed in tumor-tissue, but not non-tumor tissue, with several proteins involved in chromatin regulation including Dnmt1, Smyd3, and Eed (**Figure 4G**).

**Figure 4.**
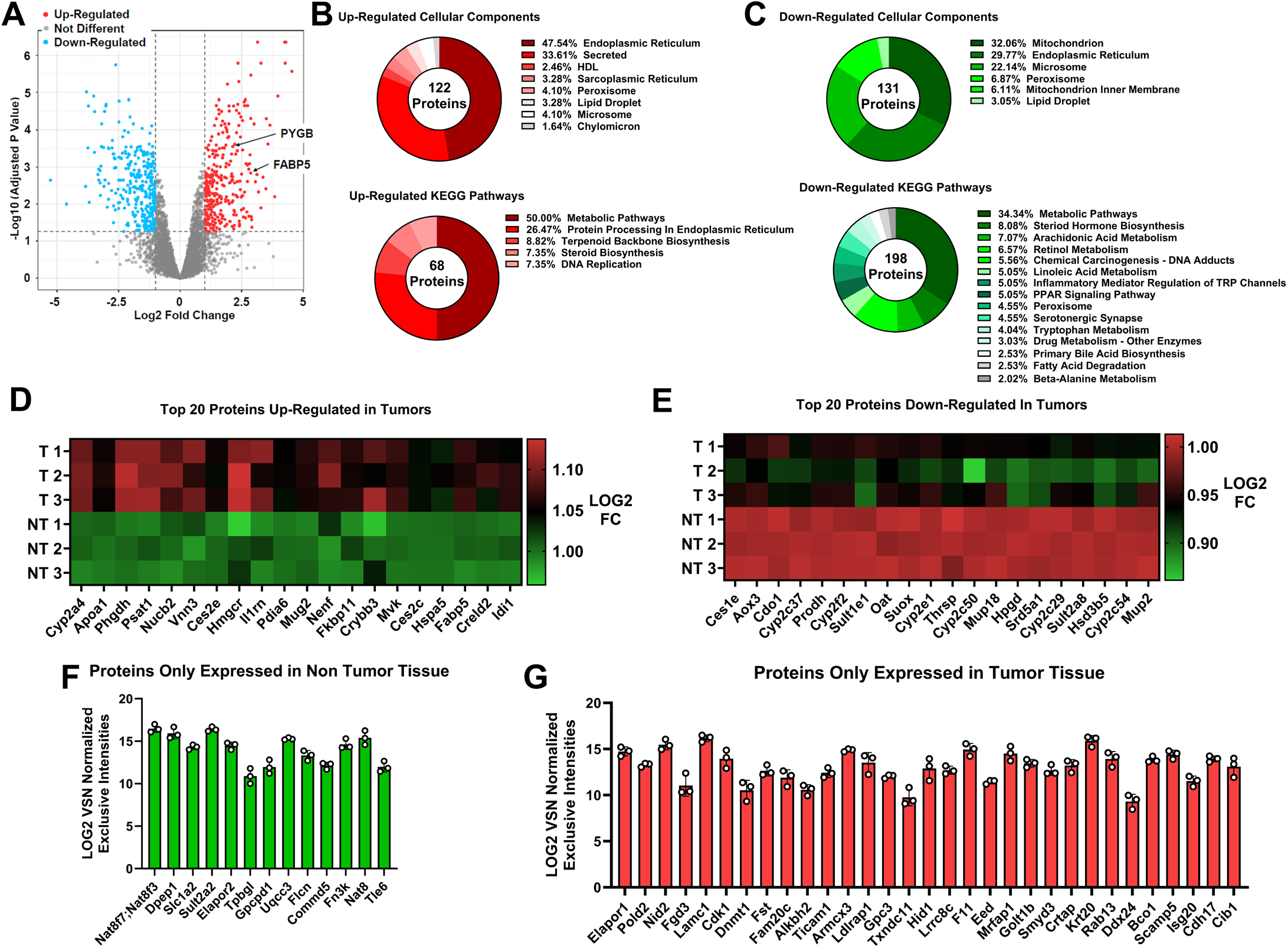
Proteomics of mouse HCC. **A)** Volcano plot illustrates significantly upregulated and downregulated proteins in tumors compared to adjacent, non-tumor tissue in WD+DEN/TAA treated mice. **B)** Select up-regulated cellular components and KEGG pathways in HCC tumors. **C)** Select down-regulated cellular components or KEGG pathways in HCC tumors. **D)** Heatmap of top 20 up-regulated proteins in tumors. **E)** Heatmap of top 20 down-regulated proteins in tumors. **F)** Proteins that were only expressed in non-tumor tissue. **G)** Proteins that were only expressed in tumors.

Some studies suggest that resting energy expenditure measured with indirect calorimetry is higher in humans with HCC compared to those without HCC (31, 32). Next, the Promethion high-resolution indirect calorimetry system was used to determine if our protocol alters whole body energy metabolism in mice. In male mice, oxygen consumption was higher at night compared to the day in the WD+DEN/TAA treated group, whereas no other significant differences existed between groups (**Figure 5A**). In female mice, oxygen consumption was highest in the WD+DEN/TAA group at night compared to the WD+DEN/TAA group during the day or compared to the LFD group during the day or night (**Figure 5B**). In male mice, the respiratory exchange ratio (RER) was higher during the night compared to the day, regardless of group (**Figure 5C**). In female mice and during the day, the RER was lower in the WD+DEN/TAA group compared to the LFD group (**Figure 5D**). During the night, these differences in RER between groups disappeared. Moreover, the RER was greater at night in both LFD and WD+DEN/TAA groups compared to the day. These differences in oxygen consumption and substrate metabolism were not associated with changes in physical activity levels (**Figure 5E-5F**) or food intake (**Figure 5G-5H**). Moreover, whole body complete glucose oxidation to CO_2_ after an I.P. injection of U^13^C glucose was not different between the groups (**Figure 5I-5L**).

**Figure 5.**
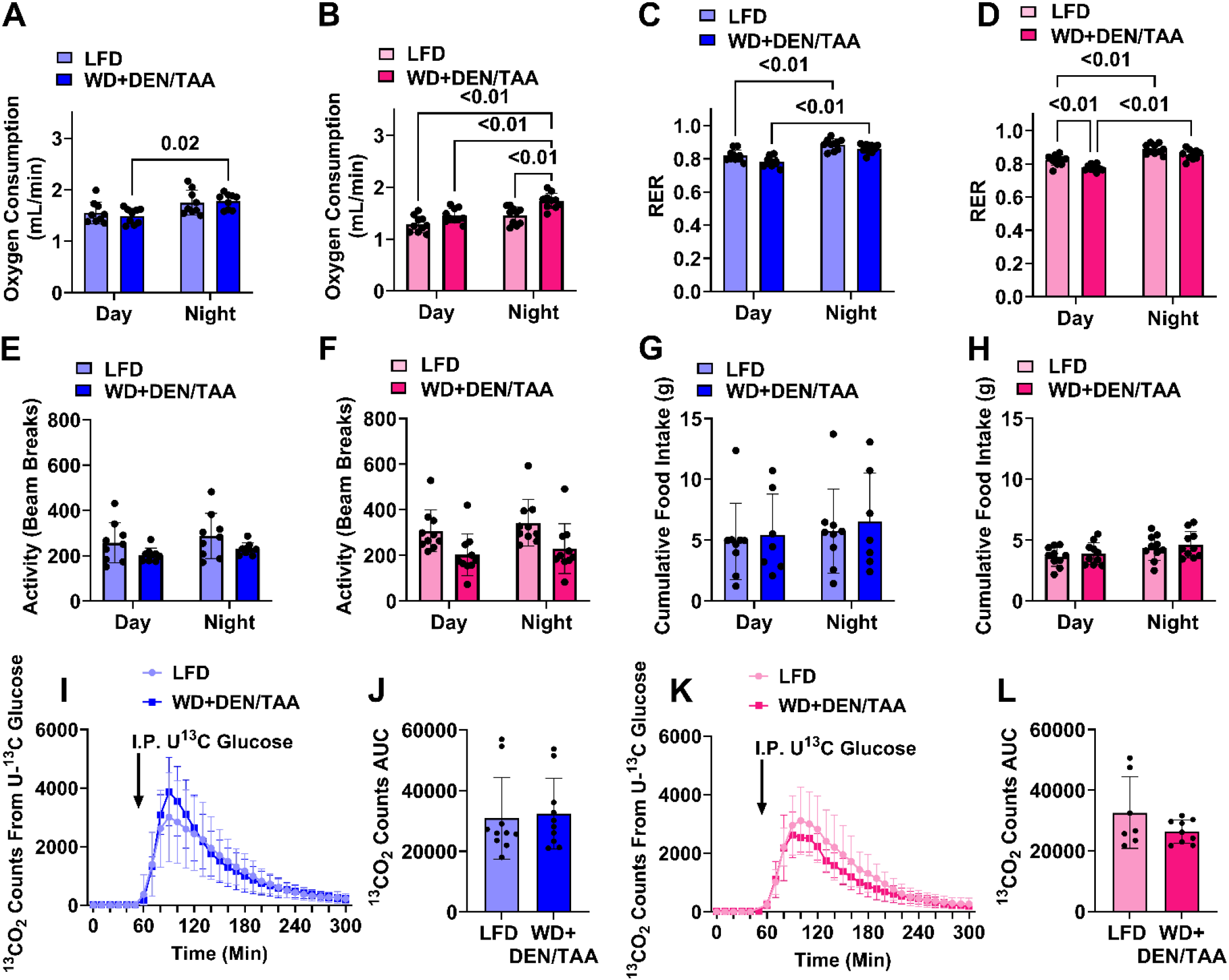
Whole body metabolism, physical activity, and food intake. **A-B)** Average oxygen consumption rates during the day and night in male and female mice. **C-D)** Average RER during the day and night. **E-F)** Average physical activity levels (beam breaks). **G-H)** Cumulative food intake during the day and night. **I-L)** Whole body glucose oxidation measured as amount of ^13^CO2 in exhaled air. Data are presented as means ± S.E.M.

## Discussion

Preclinical mouse models that closely mimic the human disease are a foundational element of translational science and are required to identify and test the efficacy and safety of new therapies prior to initiating human clinical trials. DEN is a very common genotoxic compound used to promote HCC in preclinical mouse models, yet its use results in inconsistent HCC occurrence, particularly in female mice. In this study, we resolved this issue and report that eight DEN injections spaced one week apart and starting at two weeks of age followed by TAA administration for four weeks, all on top of a Western diet, consistently and reliably promote advanced stage 2-3 HCC in both male and female mice by 30 weeks of age. Importantly, the protocol altered serum or liver HCC markers in a direction that is consistent with advanced human HCC, making this preclinical mouse model highly translational. This knowledge will be useful for future studies examining the pathogenesis of HCC or testing new therapeutic approaches for advanced HCC.

Lipid metabolism undergoes a significant remodeling in the liver cancer microenvironment, moving toward a state of increased lipid synthesis, uptake, and storage to fuel rapid cell growth, proliferation, and survival in the stressful tumor microenvironment. Our spatial transcriptomics analysis in conjunction with proteomics and western blotting revealed that FABP5, a lipid chaperone protein involved in lipid trafficking and membrane synthesis, was upregulated in tumors compared to adjacent, non-tumor tissue in our mouse model, and this effect mirrored what was observed in our human HCC samples. Previous studies have shown that FABP5 is upregulated in human advanced HCC and associated with poor prognosis and lower survival rates (27, 33, 34). In cell culture, upregulating FABP5 increases proliferation (27, 33), while knockdown of FABP5 reduces proliferation in various HCC cell lines (27). Moreover, in mice genetic ablation of FABP5 reduced HCC burden (35). Taken together, FABP5 is a physiological marker of human HCC that is also upregulated in our preclinical mouse model.

Glycogen phosphorylase brain isoform (PYGB) is one of three isoforms of glycogen phosphorylase and facilitates glycogenolysis to form glucose 1-phosphate or glucosamine 1-phosphate (36). Although initially found to be highly expressed in brain tissue, its expression is high in other tissues and various cancers (37), including HCC (34, 38). In HCC, PYGB is a better predictor of poor prognosis compared to glycogen phosphorylase liver isoform (PYGL) (38), while its inhibition reduces cell proliferation in HCC cell models (38, 39). Although localized to the cytoplasm, PYGB is also found in the nucleus where it promotes glycogenolysis to provide substrates for histone acetylation (37). The mouse model presented herein upregulates PYGB similar to human HCC, indicating that our model recapitulates a key feature of human HCC. mTOR is an atypical serine/threonine kinase and a master growth regulator that is recruited to the lysosome prior to activation. mTOR responds to a diverse range of extracellular and intracellular cues to control cell growth and limit catabolism (40). mTOR signaling is upregulated in human HCC (41, 42), while in a mouse model overactivation of mTOR is sufficient to drive HCC development (43). Consistent with this, mTOR signaling was elevated in our mouse model. These findings indicate that our model will be suitable to test how mTOR inhibitors impact HCC.

In conclusion, this study introduces, for the first time, a reliable chemical carcinogen model that induces advanced HCC in both male and female mice. Importantly, the model was highly tolerable and induced protein and gene signatures comparable to human HCC. This new protocol will be a valuable model for preclinical testing of new therapeutic approaches for advanced stage HCC.

## List of Abbreviations

ALT: alanine aminotransferase
AST: aspartate aminotransferase
DEN: diethylnitrosamine
DNA: deoxyribonucleic acid
FABP5: fatty acid binding protein 5
FFPE: formalin fixed paraffin embedded
HCC: hepatocellular carcinoma
mTOR: mechanistic target of rapamycin
MUP20: major urinary protein 20
LFD: low-fat diet
PYGB: glycogen phosphorylase brain isoform
TAA: thioacetamide
WD: Western diet

## Acknowledgements

We would like to thank Shirley Ennis and Stephanie Wong for helping with tissue embedding, sectioning, staining, and imaging. This study was financially supported by National Institute of Health grants R35GM154665 and P20GM135002 (subprojects 8000, 8438, and 5359) awarded to TDH. This project used the Cell Biology and Bioimaging Core as well as the Genomics core at Pennington Biomedical Research Center; both Cores are supported in part by National Institute of Health awards P20GM135002 and P30DK072476. The proteomics experiment was supported by an IDeA National Resource for Quantitative Proteomics National Institute of Health grant R24GM137786. BioRender was used to create the schematic figures used in the manuscript.

## Author Contributions

MCM, ERA, TDH; conception and design, data acquisition, data analysis, data interpretation, drafting article, final approval. DHB, RC, SW, JS, JMS, SGM: data acquisition, data analysis, data interpretation, article revision, final approval.

